# Fully Phased Telomere-to-Telomere Assemblies for Thoroughbred Horse and Donkey Haplotypes derived from a Mule Illuminate the Peculiar Evolution of Equid Centromeres

**DOI:** 10.64898/2026.02.26.707634

**Authors:** Kai Li, Eleonora Cappelletti, Carey Dessaix, Julia Ciosek, Emily Robyn, Lauren Johnson, Nahla Hussien AbouEl Ela, Xiomara Arias, David L. Adelson, Terje Raudsepp, Francesca M. Piras, Melissa Laird-Smith, Elizabeth Hudson, Brandon D. Pickett, Sergey Koren, Brian P. Walenz, Shelise Y. Brooks, Christina Sison, Juyun Crawford, Gerard Bouffard, Adam M. Phillippy, Donald Miller, Douglas F. Antczak, Jonah Cullen, Sam Stroupe, Brian Davis, Molly McCue, Sian Durward-Akhurst, Jessica L. Petersen, Elena Giulotto, Ted Kalbfleisch

## Abstract

We present telomere-to-telomere genome assemblies of a Thoroughbred horse and a donkey derived from their mule offspring. Now adopted and annotated by NCBI as reference genomes, these assemblies resolve previously inaccessible regions, including satellite arrays, duplications, and telomeres. Equids are known to exhibit an uncoupling between satellite DNA and centromeric function. The completeness of these assemblies enabled annotation of both satellite-based and satellite-free centromeres, as well as non-centromeric satellite loci, revealing notable centromeric plasticity. They also allowed detailed characterization of the variable binding domains of CENP-A—the epigenetic determinant of centromere identity—and CENP-B, whose association with CENP-A, previously considered typical based on a few model organisms, is absent in equids. Comparative analyses of satellite repeats and centromere positions provide new insights into the accelerated karyotypic reshuffling in equid evolution. These assemblies represent foundational resources for equid genomics and support ongoing initiatives such as the Equine Pangenome Project.

## Background

Genome resources for the horse have advanced at more or less the same rate as those of other large animal agricultural species such as cattle, pigs, and sheep. The first reference genome for the horse was released in 2007 and referred to as EquCab2.0[1] (GCF_000002305.2). It was followed in 2018 by EquCab3.0[2] (GCF_002863925.1), which was improved in both composition, and contiguity by high throughput sequencing technologies that had recently become available. Those assemblies were foundational to the hundreds of horse genetic and genomic studies that followed including the Equine FAANG project, and made possible other high throughput tools including the high density SNP chips: 50k in 2010, 70k[3] in 2012, and the 670k[4] in 2017. As sequencing technologies and the tools to assemble these data have dramatically improved in terms of both length and accuracy, our efforts to build ever better assemblies that are more contiguous, complete, and accurate have kept pace.

By virtue of both long, and ultra-long read technologies, we are now able to uniquely resolve repeat regions and characterize centromeres. Centromeres are essential nucleoprotein structures which ensure chromosome segregation during cell division. Although centromeres are discrete genetic loci, their function is not determined by the underlying DNA sequence but epigenetically specified by the histone H3 variant CENP-A[5]. While centromeric DNA sequences are neither sufficient nor required to specify centromere function, mammalian centromeres are typically associated with extended arrays of satellite DNA[6], which remained largely absent from earlier genome assemblies. This has posed a significant challenge in unraveling centromeric function at the molecular level. In this context, a turning point was the discovery that equid species are characterized by the coexistence of satellite-based and satellite-free centromeres [1,7–10]. The presence of functional satellite-free centromeres made the genus *Equus* a powerful model system in centromere biology[11–17]. In the horse (2n = 64), only the centromere of chromosome 11 is satellite-free while the other centromeres are satellite-based [1,8]. In the donkey (2n = 62), 16 centromeres are satellite-free, demonstrating that although more than half of centromeres are devoid of satellite DNA, this configuration is compatible with genome stability and species survival[8]. The positions of these satellite-free centromeres, identified as CENP-A binding domains, are not fixed but slide in an about 600 kb region, giving rise to “epialleles” that follow Mendelian inheritance[8,18]. However, at several centromeres, substantial centromere movements, on the order of 50-80 kb, were observed from parent to offspring, indicating that the position of CENP-A binding domains can slide in one generation[8]. In contrast, the position of the centromere is stable during mitotic propagation of cultured cells[8] and during tissue differentiation[16], suggesting that centromere sliding may occur during meiosis.

The genus *Equus* remains unique among mammals for its abundance of satellite-free centromeres[1,7–9], which is attributed to the rapid and recent evolution of equid species. In a very short evolutionary time frame, frequent centromere repositioning events and chromosomal rearrangements, particularly centric fusions, reshaped equid karyotypes [7–10,17,19,20]. These mechanisms gave rise to the numerous satellite-free centromeres which are considered “immature” because they did not have sufficient time to acquire the structural and sequence complexity typical of mammalian centromeres. Another consequence of this great karyotype reshuffling during equid evolution was the heterogeneous organization of satellite DNA sequences[7,17]. Several satellite families were described in equids, namely 37cen, 2PI, CENPB-sat, EC137, SatA, SatB, SatC, SatD and SatE[7,17,21,22]. These loci are present either at satellite-based centromeres or at non-centromeric positions, corresponding to ancestral inactivated centromeres[7,17]. Recent findings revealed that CENPB-sat, which is the satellite bound by the centromeric protein CENP-B, is not the major centromeric satellite in any equid species[17]. This discovery challenges the common view, mainly based on human and mouse experimental systems, that CENP-B, the unique centromeric protein binding a specific motif (CENP-B box)[23], binds the main centromeric satellite and thus increases centromere strength and segregation fidelity[24]. In equids, CENP-B is expressed but does not bind the majority of satellite-based or satellite-free centromeres, while it is localized at several non-centromeric terminal positions, likely corresponding to ancestral, now-inactive, centromeres[17]. However, centromeres lacking CENP-B are functional and do not show any segregation defect in mitosis compared to CENP-B positive chromosomes[17]. According to our recent model, the equid CENPB-sat satellite was centromeric in the equid ancestor, lost centromeric function during evolution, and gave rise to a shorter CENP-A bound repeat (37cen) not containing the CENP-B box[17]. Over evolutionary timescales, this uncoupling between CENP-B and CENP-A may have played a role in the extensive evolutionary reshuffling of equid centromeres.

While equid satellite-free centromeres were well characterized at the sequence level, satellite-based centromeres and satellite loci were not included in the previous, non-T2T equid genome assemblies.

Here, we present the results of our effort to create complete assemblies for horse (*Equus caballus* GCF_041296265.1) and donkey (*Equus asinus* GCF_041296235.1) haplotypes derived from a female mule named TB-T2T and EquAss-T2T_v2, respectively. The mule, a cross-species hybrid, offers an advantage for genome assembly, an approach that has already been applied successfully in cattle, bison, and yak[25–27]. Because the mule’s maternal and paternal chromosomes are from species diverged roughly four million years ago[28,29], the resulting species-specific alleles are abundant enough to reliably assign sequencing reads to their parent of origin—either before assembly using parental sequence, or during assembly as the algorithm separates and reconstructs each parental haplotype readily based on their distinct evolutionary differences. These assemblies have been adopted as the reference genomes for the two species and were subsequently annotated by NCBI in October 2024. These new T2T assemblies are essential to the ongoing annotation efforts for variation across all popular breeds, and immune response genes. A tissue repository has been created with tissues collected from the mule used in this work, and may be used to continue the annotation of this resource. In this manuscript we provide a comprehensive annotation of repetitive elements, including satellite arrays and transposons, and a characterization of centromeric domains. Comparative analyses of satellite repeats and centromere positions in horse and donkey illuminate the accelerated karyotypic reshuffling that has shaped equid evolution. Additionally, we provide an equid-focused public resource for these “community expert created” annotations that can be downloaded, or otherwise used in software or browsers capable of accessing and using web based datasets.

## Results

### Karyotype

The mule was karyotypically normal (**Supplementary Figure 1**). All twenty cells that were analyzed, had a 63,XX female hybrid karyotype composed of a normal haploid set of horse (31,X) and normal haploid set of donkey (30,X) chromosomes. Because not all horse (2n=64) and donkey (2n=62) chromosomes share one-to-one homology, the horse and donkey haploid karyotypes were arranged separately.

### Genome Composition

Both the horse (GCF_041296265.1) and the donkey (GCF_041296235.1) assemblies have been deposited to NCBI, and have been annotated as the reference genomes for their respective species. Full details of the assembled chromosomes including the numbers of telomeres and gaps (and descriptions thereof) are presented in **Supplementary Table 1**. Briefly, the horse TB-T2T assembly is a 31 chromosome + chrX and the mitochondrial (chrMT) haplotype. Both telomeres were captured on 26 chromosomes while the p-terminal end was missing at 6 chromosomes (1, 18, 19, 20, 22, and 31) (**Supplementary Table 1**). The terminal regions of the five acrocentric chromosomes were identified in five unincorporated contigs containing the missing telomeres, pericentromeric and centromeric satellite arrays described below in **Table 1A**. These were initially unplaced by the assembly pipeline but were later associated with their corresponding chromosomes. The donkey EquAss-T2T_v2 haplotype contains 30 chromosomes + chrX. Both telomeres were captured on 19 chromosomes (**Supplementary Table 1**). Seven of the missing telomeres were captured in unincorporated contigs, but not anchored to chromosomes (**Table 1B**).

**Table 1:** Unincorporated contigs in the reference assembly identified as extending chromosomal contigs in the A) TB-T2T reference genome assembly that include the missing centromeres and telomeres, and B) EquAss-T2T_v2 reference genome assembly.

| TB-T2T Chromosome | RefSeq Accession | Extending Contig | Extending Contig length | NCBI Annotated genes on extension | Contig Orientation |
| --- | --- | --- | --- | --- | --- |
| ECA chr18p | NC_091701.1 | NW_027222394.1 | 12,873,522 | 81 | - |
| ECA chr19p | NC_091702.1 | NW_027222395.1 | 13,474,092 | 70 | - |
| ECA chr20p | NC_091703.1 | NW_027222392.1 | 7,718,480 | 39 | - |
| ECA chr22p | NC_091705.1 | NW_027222391.1 | 13,048,731 | 85 | + |
| ECA chr31p | NC_091714.1 | NW_027222343.1 | 5,541,805 | 4 | - |

| EquAss-T2T_v2 Chromosome | RefSeq Accession | Extending Contig | Extending Contig length | NCBI Annotated genes on extension | Contig Orientation |
| --- | --- | --- | --- | --- | --- |
| EAS chr1p | NC_091790.1 | NW_027224856.1 | 569,576 | 18 | + |
| EAS chr19q | NC_091808.1 | NW_027225534.1 | 9,345,831 | 1 | - |
| EAS chr22p | NC_091811.1 | NW_027224843.1 | 1,941,500 | 8 | + |
| EAS chr25p | NC_091814.1 | NW_027225523.1 | 10,933,035 | 0 | + |
| EAS chr26p | NC_091815.1 | NW_027225316.1 | 523,695 | 3 | - |
| EAS chr28p | NC_091817.1 | NW_027225530.1 | 642,182 | 3 | - |
| EAS chr30q | NC_091819.1 | NW_027224899.1 | 423,658 | 0 | + |

Comparing the new TB-T2T assembly to EquCab3.0 demonstrates that improvements to nearly all chromosomes were made to the centromeric regions (**Figure 1**). Assemblies attempted either with PacBio HiFi (HiFi) only, or Oxford Nanopore Technologies (ONT) reads only are not able to span these regions gaplessly. Even with the improvements typical of HiFi + ONT assembly efforts, five gaps remain in the otherwise contiguous chromosomes that could not be resolved in the horse. They are located on TB-T2T chromosomes 16, 19, and 27 (**Supplementary Table 1**).

**Figure 1:**
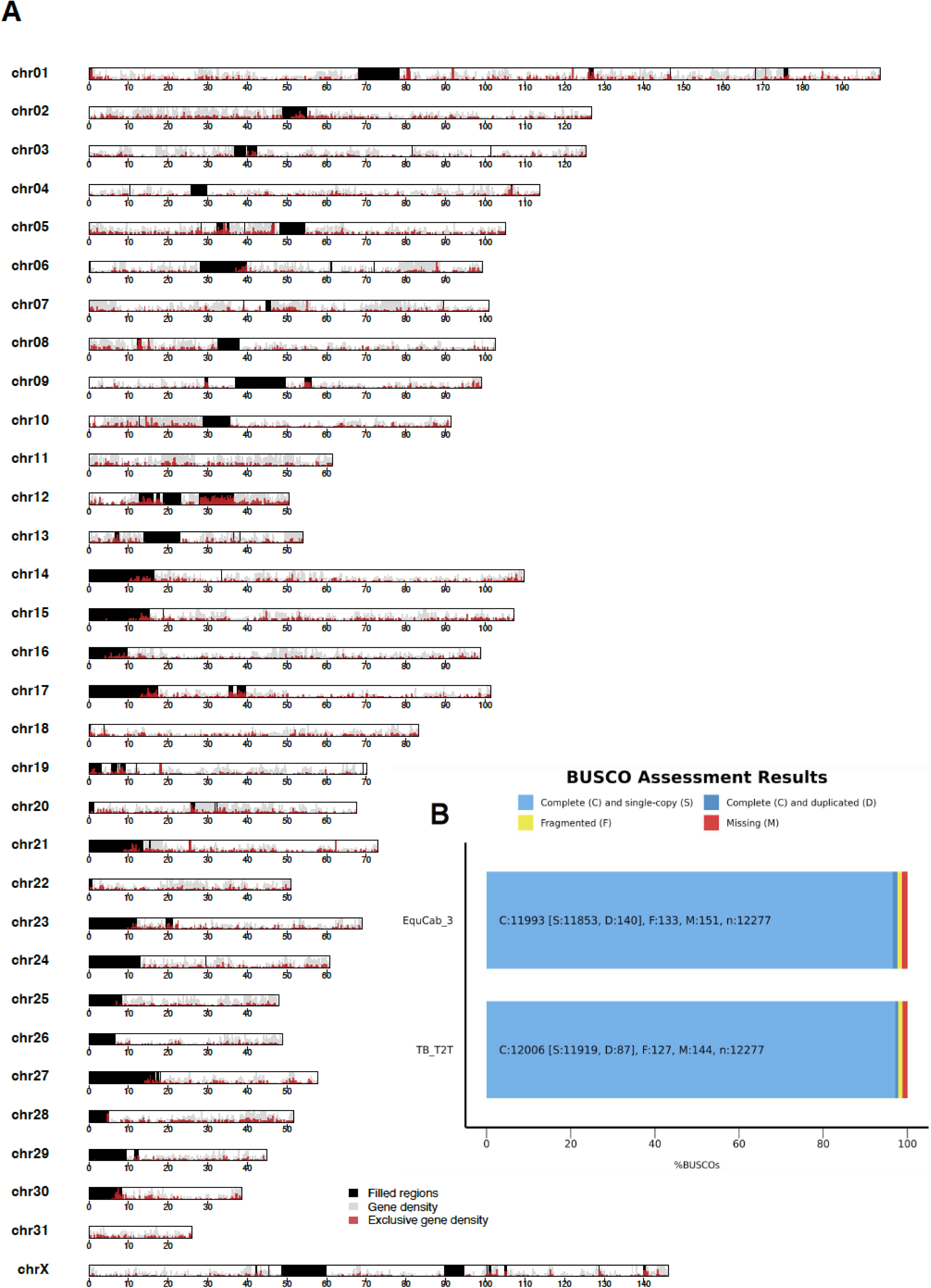
A summary of the TB-T2T genome assembly. (A) An ideogram of TB-T2T assembly features. For each chromosome (chr), the following information is provided: the density of genes found in the TB-T2T, and not EquCab3 is shown in red, the density of genes reported previously in EquCab3.0 are in gray, and gaps and issues in EquCab3.0 fixed by TB-T2T are shown in black. (B) BUSCO completeness analysis based on the mammalia_odb12 dataset showing the proportions of complete (C; single-copy [S] and duplicated [D]), fragmented (F), and missing (M) orthologs. The horse telomere-to-telomere (T2T) assembly TB-T2T exhibits slightly higher completeness and fewer missing BUSCOs compared to the corresponding EquCab3 reference assembly, indicating improved assembly quality and gene representation.

### BUSCO and Merqury Analyses

Benchmarking Universal Single-Copy Orthologs (BUSCO) and Merqury are tools that assess completeness biologically, and compositionally respectively. BUSCO identifies genes conserved across mammalian species, whereas Merqury identifies sequence elements found in the source sequence data (ONT, HiFi, and Illumina short reads) from the source animal that are not present in the assembly they produced, and vice versa.

The BUSCO analysis revealed a clear improvement in assembly completeness at the gene level. The number of missing conserved mammalian genes decreased from 150 to 143 in the horse assembly (**Figure 1B**) and from 177 to 140 in the donkey assembly (**Figure 2B**). Notably, the number of missing and fragmented BUSCOs was reduced in the T2T assemblies, indicating successful resolution of previously incomplete or duplicated regions. These results highlight that our assemblies captured more of the conserved mammalian gene content, reflecting improved contiguity and completeness achieved using the T2T assembly strategy.

**Figure 2:**
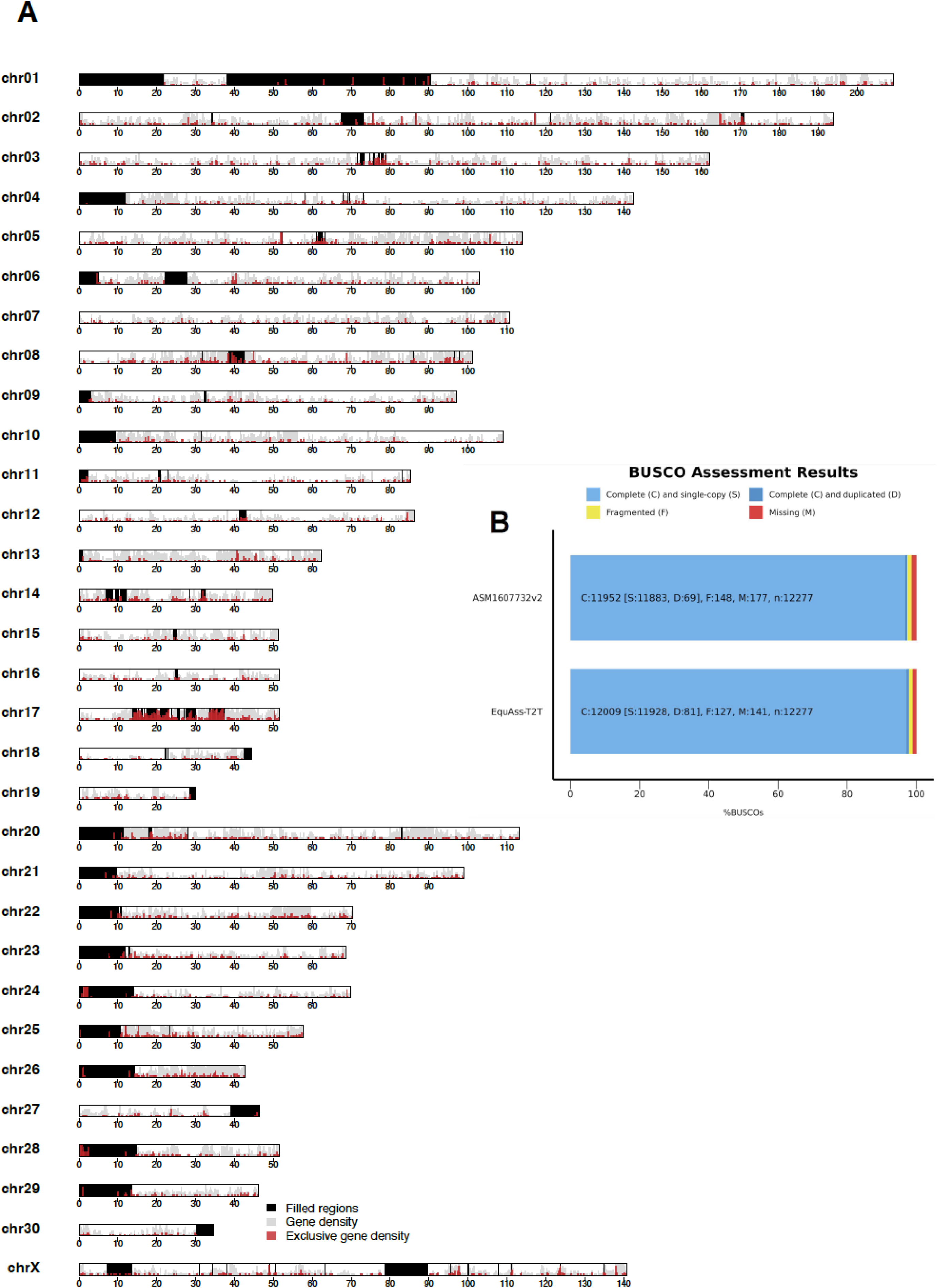
A summary of the EquAss-T2T_v2 genome assembly. (A) An ideogram of the EquAss-T2T_v2 assembly features. For each chromosome (chr), the following information is provided: the density of genes found in the EquAss-T2T_v2 and not ASM1607732v2 is shown in red, the density of genes reported previously in ASM1607732v2 are in gray, and gaps and issues in ASM1607732v2 fixed by EquAss-T2T_v2 are shown in black. B)BUSCO completeness analysis based on the mammalia_odb12 dataset showing the proportions of complete (C; single-copy [S] and duplicated [D]), fragmented (F), and missing (M) orthologs. The donkey telomere-to-telomere (T2T) assembly EquAss-T2T_v2 exhibits slightly higher completeness and fewer missing BUSCOs compared to the corresponding ASM1607732v2 reference assembly, indicating improved assembly quality and gene representation.

Merqury is a software package that evaluates the quality and completeness of the assembly by comparing a set of k-mers derived from high-accuracy sequence reads to a genome assembly[30]. The Merqury analysis produced a quality score of 52.5. **Figure 3A** reveals 2,682,580,525 and 2,694,786,446 kmers that are found in the TB-T2T and EquAss-T2T_v2 respectively. The graph is consistent with a kmer profile of an interspecies cross with larger haplotype specific peaks at roughly half coverage of the diploid genome.

**Figure 3:**
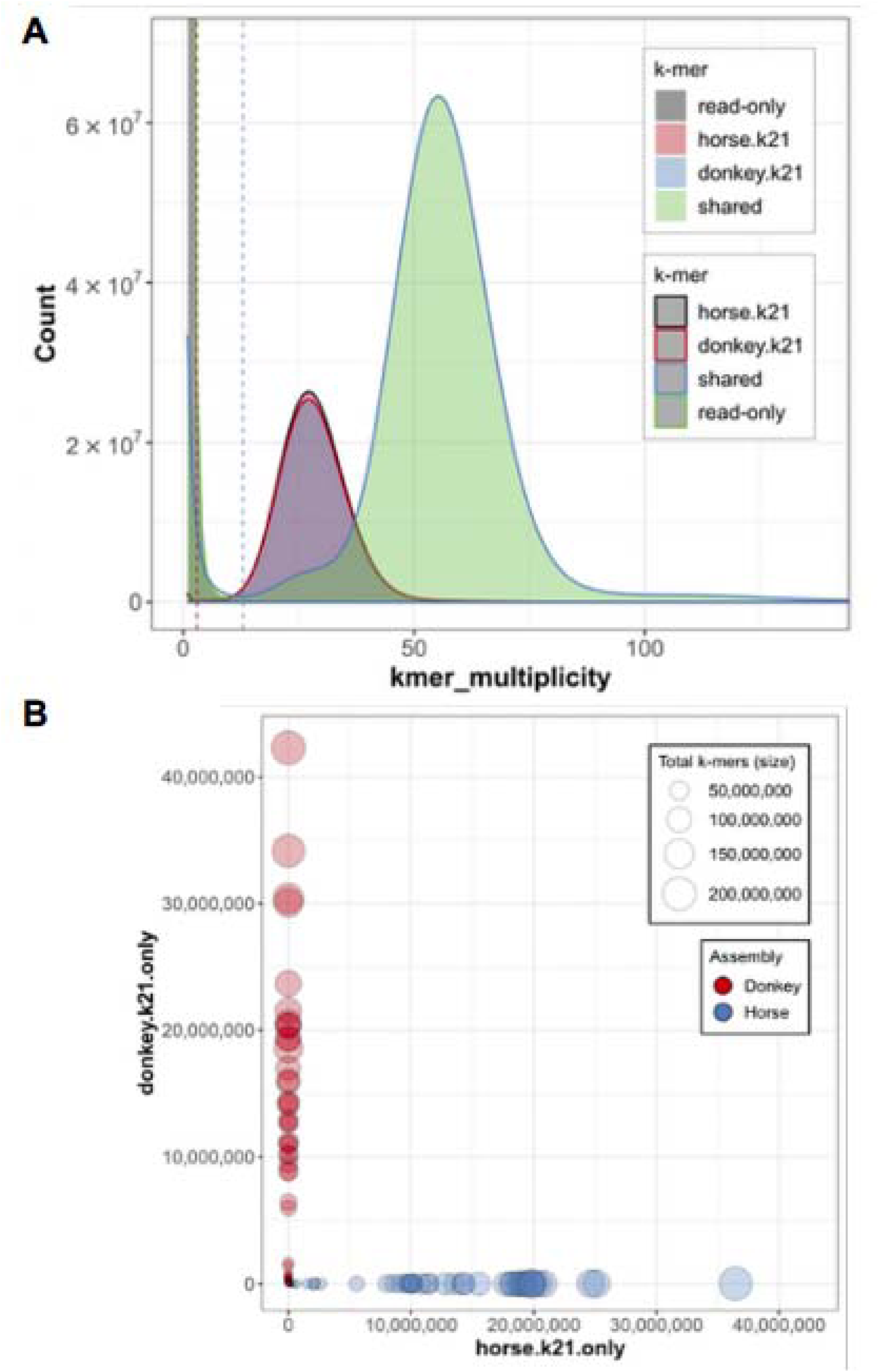
k-mer–based assembly evaluation and haplotype assignment of the mule genome. (a) Merqury k-mer spectrum analysis comparing donkey, horse, and shared k-mers demonstrates high completeness and consistency of the mule assembly. The majority of k-mers are shared (green), indicating accurate sequence representation with minimal read-only regions. (b) Haplotype-specific k-mer plot showing parental assignment accuracy in the mule assembly. Each point represents a contig, with red and blue denoting donkey- and horse-specific k-mers, respectively. The clear separation of donkey contigs along the Y-axis and horse contigs along the X-axis indicates near-perfect phasing and minimal cross-haplotype contamination, confirming the faithful resolution of both parental haplotypes.

The clear segregation of points along the axes—donkey contigs clustering on the Y-axis and horse contigs on the X-axis—demonstrates that each haplotype was cleanly resolved with minimal cross-contamination (**Figure 3B**). The absence of mixed k-mers (points near the diagonal) highlights the near-perfect phasing, confirming that the assembly faithfully distinguishes the two parental haplotypes with exceptional accuracy.

### Satellite DNA and centromere organization in the horse

In previous work, we demonstrated that, in the horse, the centromere of chromosome 11 is completely devoid of satellite DNA, whereas all other centromeres are satellite-based [1,7,8,15]. While this satellite-free centromere has been thoroughly characterized at the molecular level, satellite-based centromeres have remained largely uncharacterized at the sequence level due to their highly repetitive nature.

To investigate the organization of satellite DNA in the horse genome, we first mapped the satellite DNA families previously identified in the horse (37cen, 2PI, CENPB-sat, EC137, SatA and SatB) onto the TB-T2T reference genome using RepeatMasker analysis. The monomer size and the consensus sequence of each satellite are reported in **Supplementary Table 2**. The genomic coordinates of the various satellite arrays are provided in **Supplementary Table 3.** As shown in **Figure 4**, satellite repeats constitute about 9% of the genome. Consistent with our previous data obtained from unassembled short reads[17], 37cen and 2PI are the most abundant satellites, while CENPB-sat, EC137, SatA and SatB are represented at lower levels (**Figure 4-A**).

**Figure 4.**
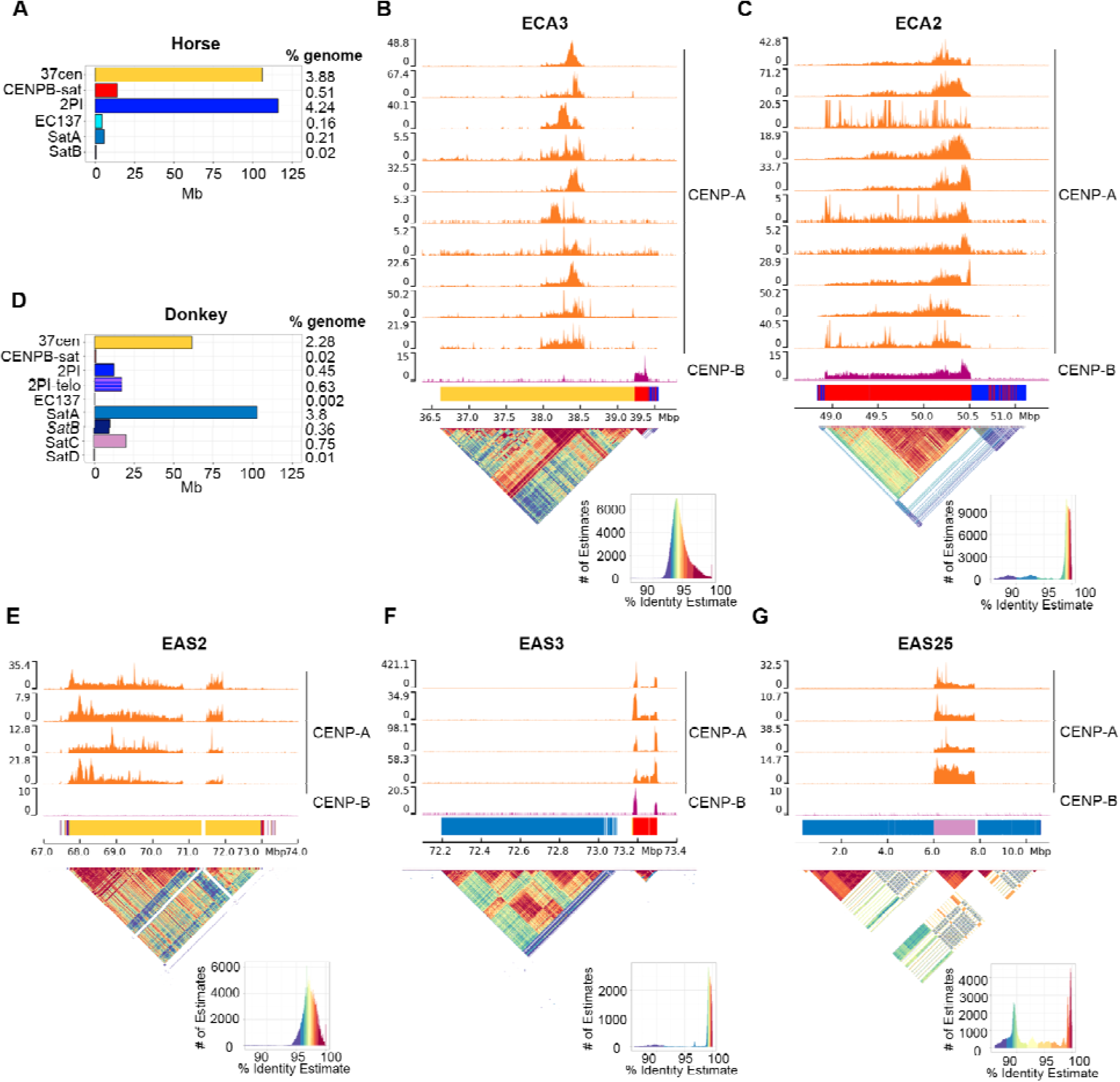
Horse and donkey satellite-based centromeres. A) Barplot of the total lengths in megabases of each satellite family in the horse TB-T2T genome. For each family, the genome percentage is reported on the right. B) Organization of the centromere of horse chromosome 3, as representative example of the satellite-based horse centromeres. The ChIP-seq profiles of CENP-A (orange, 10 individuals) and CENP-B (purple, 1 individual) are shown at the top. The y-axis reports the normalized read counts whereas the x-axis reports the coordinates on the horse reference genome. The colored bar represents the satellite DNA array, with colors denoting different satellite families as indicated in panel A. The sequence identity maps obtained with ModDotPlot are reported at the bottom. C) The satellite-based centromere of horse chromosome 2. This is the only horse centromere where CENP-A and CENP-B bind the same satellite repeat. D) Barplot of the total lengths in megabases of each satellite family in the donkey EquAss-T2T_v2 genome. For each family, the genome percentage is reported on the right. E) The satellite-based centromere of donkey chromosome 2. F) The satellite-based centromere of donkey chromosome 3. G) The satellite-based centromere of donkey chromosome 25. Panels C, E, F and G are structured as panel B. In panels B and C, the CENP-B individual is the 7th from the top within the CENP-A tracks; in E–G, it is the 2nd from the top within the CENP-A tracks.

Following the annotation of satellite repeats in the TB-T2T assembly, we analyzed CENP-A ChIP-seq datasets from ten horse individuals to investigate the organization of centromeric domains. Nine of these datasets were generated in our previous studies[8,16,17], while one was produced specifically for this study. In addition, we examined the relative positioning of CENP-A and CENP-B binding domains using a CENP-B ChIP-seq dataset from one of these individuals [17]. The results of this comprehensive analysis of satellite DNA and centromere organization are shown in **Figure 4-B-C** and **Supplementary Figures: 2, 3, 4**.

We confirmed that, in the horse, only the chromosome 11 (ECA11) centromere is satellite-free (**Supplementary Figure 2**). At this locus, the profiles of CENP-A enrichment peaks across different horse individuals were markedly improved compared to those obtained using the previous EquCab3.0 reference genome[16], further supporting the high quality of the TB-T2T assembly. As previously shown using different reference genomes[16,18], the position of CENP-A enrichment peaks varies in the different individuals within a 500 kb window, confirming the presence of epialleles for CENP-A binding in the population[8,18]. In agreement with our recent findings[17], no CENP-B binding is present at this satellite-free centromere (**Supplementary Figure 2**).

With the exception of the ECA11 centromere, all other centromeres are associated with satellite DNA (**Figure 4*-*B-C**, **Supplementary Figure 3**). An example of a typical horse satellite-based centromere is that of ECA3 (**Figure 4*-*B)**. This chromosome comprises an extended array of 37cen which is flanked by shorter arrays of CENPB-sat and 2PI satellites. The mapping of CENP-A ChIP-seq datasets revealed that CENP-A enrichment peaks from the 10 analyzed horse individuals are not distributed across the entire 37cen array. Instead, they are confined to a central 500 kb region, which exhibits the highest sequence identity according to the identity heatmap generated by ModDotPlot, and is embedded within more divergent repeat units at the edges. Interestingly, the position of the CENP-A binding domain varies among the different individuals, similar to what we previously discovered for the horse satellite-free centromere of chromosome 11[16,18]. This finding indicates that the functional centromere is located in the most sequence-conserved portion of the 37cen array, further confirming that 37cen is the satellite bearing the centromeric function in the horse[17,22], while the remaining portion of the array is pericentromeric. CENPB-sat and 2PI satellites are not bound by CENP-A and are located in the peripheral region of the pericentromere. At this centromere, CENP-B binding was detected at the short 180 kb CENPB-sat array, while it is absent at the centromeric domain confirming the dissociation between CENP-A and CENP-B domains at equid centromeres[17]. This centromere organization is conserved across most equid satellite-based centromeres, with the CENP-A binding domain localized within the most conserved regions of the 37cen arrays (**Supplementary Figure 3**). In different cases, such as ECA6, ECA7, ECA10, ECA12, ECA14, ECA15 and ECA26, variation in the CENP-A binding position among individuals has been observed (**Supplementary Figure 3**).

The metacentric chromosomes show long arrays of 37cen flanked by short arrays of CENPB-sat, 2PI and EC137 (**Supplementary Figure 3**). Differently, acrocentric chromosomes are often characterized by shorter arrays of 37cen and longer arrays of CENPB-sat and 2PI compared to metacentric ones (**Supplementary Figure 3**). Interestingly, these satellite arrays lie directly adjacent to the (TTAGGG)n telomeric repeats (**Supplementary Figure 3**). Consistent with our recent findings[17], the ECA11 satellite-free centromere and numerous horse satellite-based centromeres (ECA1, ECA4, ECA5, ECA7, ECA9, ECA12, ECA13, ECA27, ECA28, ECA29 and ECAX) contain little to no CENPB-sat arrays and do not exhibit CENP-B binding (**Supplementary Figures 2, and 3**). At the other satellite-based centromeres, CENPB-sat arrays are pericentromeric and often intermingled with 2PI repeats (**Supplementary Figure 3**), confirming the dissociation between CENP-A and CENP-B domains[17].

The only exception to this organization is the centromere of ECA2 (**Figure 4-C**). We previously proposed that this centromere, lacking 37cen repeats, retains the ancestral configuration of *Equus* centromeres, with CENPB-sat binding both CENP-A and CENP-B[17]. In agreement with this hypothesis, this centromere is characterized by an extended 1.5 Mb CENPB-sat array which is bound by both CENP-A and CENP-B proteins (**Figure 4-C**). This array is flanked by degenerated arrays of 2PI intermingled with very short and degenerated CENPB-sat repetitions, while no 37cen is present.

In the primary assembly, no CENP-A enrichment peaks were detected at the ECA18, ECA19, ECA20, ECA22 and ECA31 chromosomes (**Supplementary Table 3**). The entire p arm of these chromosomes, encompassing the telomeric end and the centromeric region, was missing. After the annotation of the assembly by NCBI, we demonstrated that these five missing ends were present in five unplaced contigs, which we were able to anchor to the corresponding chromosomes (**Table 1A**). These sequences, which include telomeric (TTAGGG)n repeats, pericentromeric and centromeric satellite arrays, were included in the analyses shown in **Supplementary Figure 3**. These five centromeres exhibit an organization comparable to that of the other acrocentric chromosomes, with satellite arrays extending from the telomeric boundary.

The presence of SatA and SatB at horse satellite-based centromeres is negligible (**Supplementary Table 3)**, with only a few kilobase-long stretches detected at ECA26 for SatA, and at ECA25, ECA28 and ECA31 for SatB (**Supplementary Figure 3**). Differently, a 5 Mb array of SatA is present at the q-arm of ECAX and represents the only extended non-centromeric satellite array of the horse genome (**Supplementary Figure 4**). At this locus, neither CENP-A or CENP-B binding was detected (**Supplementary Figure 4**). It is important to note that a RepeatMasker analysis showed that SatA units contain regions of identity with LINE L1-4B elements, suggesting that these retrotransposons may be involved in the expansion of SatA arrays. Short stretches of satellite repeats, ranging from a few base pairs to a few kilobases, are otherwise present in other genomic positions, mainly within a few megabases of the centromeric and pericentromeric satellite arrays (**Supplementary Table 3**). Interestingly, quite an extended (370 kb) array of SatB is present at the p-terminus of ECA1 (**Supplementary Table 3**) where it could represent a relic of an ancestral satellite-based centromere.

### Satellite DNA and centromere organization in the donkey

Our previous work demonstrated that, in the donkey, 16 out of 31 centromeres were satellite-free and coexist with satellite-based centromeres, alongside several non-centromeric satellite loci that represent relics of ancestral centromeres inactivated during evolution[7,8].

To characterize the organization of satellite DNA in the donkey genome, we first localized the different satellite DNA families of this species (37cen, 2PI, CENPB-sat, EC137, SatA, SatB, SatC, SatD) onto the EquAss-T2T_v2 reference genome using RepeatMasker analysis. The monomer size and the consensus sequence of each satellite are reported in **Supplementary Table 2**. The genomic coordinates of the various satellite arrays are provided in **Supplementary Table 4**. As shown in **Figure 4-D**, satellite repeats constitute about 8.3% of the genome. Consistent with our previous data obtained from unassembled short reads[17], 37cen and SatA are the most abundant satellites, while 2PI, SatB and SatC are less prevalent, and CENPB-sat, EC137 and SatD are nearly absent (**Figure 4-D**). Interestingly, we discovered that the majority of 2PI loci are composed by an undescribed subfamily of about 1.2 kb monomers, consisting of about 220 bp of telomeric-like repeats followed by 2PI repeats (**Supplementary Figure 5**). This subfamily, termed 2PI-telo, accounts for more than half of the total 2PI satellite content (**Figure 4-D**).

Following the annotation of satellite repeats in the EquAss-T2T_v2 assembly, we analyzed CENP-A ChIP-seq datasets from four donkey individuals to investigate the organization of centromeric domains. Two of these datasets were generated in our previous work[8], while the other two were produced specifically for this study. In addition, we used a CENP-B ChIP-seq dataset, which was previously obtained from one of these individuals[17], to inspect the relative positioning of CENP-A and CENP-B binding domains. The comprehensive characterization of satellite DNA and centromere organization is presented below and illustrated in **Figure 4-E-G** and **Supplementary Figures 6, 7, 8 and 9**.

The mapping of the CENP-A ChIP-seq datasets from four individuals confirmed that 16 centromeres are satellite-free (**Supplementary Figures 6, and 7**). The CENP-A enrichment peaks generally display a regular Gaussian-like shape, supporting the high quality of this assembly. Irregular peaks are present at the centromeres of *Equus asinus* chromosomes 13 (EAS13) and EAS14, suggesting that these loci may be rearranged in the donkey individuals analyzed by ChIP-seq compared to the reference genome. We previously showed that five satellite-free centromeres contain DNA duplications[8]. This sequence organization was confirmed by the identity heatmaps obtained with ModDotPlot (**Supplementary Figure 7**). This repetitive organization was particularly evident at the EAS8 centromere, where duplications span approximately 4 Mb. In contrast, at EAS19, the repeated organization was not detectable by ModDotPlot in agreement with the previously described polymorphism in the population regarding the presence of duplications[8]. At all satellite-free centromeres, the position of CENP-A enrichment peaks varies in the four different individuals (**Supplementary Figure 6, and 7**), confirming the presence of epialleles for CENP-A binding in the population[8,18]. In our previous work we proved that CENP-A binding domains can move within regions of up to 600 kb[8]. However, the analysis of these four individuals revealed that, at the EAS12 satellite-free centromere, CENP-A binding domains are distributed within a 2.8 Mb window, with Donkey D exhibiting two epialleles separated by 2.5 Mb (**Supplementary Figure 6**). This finding demonstrates that centromere sliding can occur over a larger genomic region than previously anticipated. In agreement with our recent findings[17], no CENP-B binding is present at any satellite-free centromere (**Supplementary Figure 6, and 7**).

The analysis of the 15 satellite-based centromeres revealed that the organization of these centromeres is heterogeneous in the donkey (**Figure 4-E-G**, **Supplementary Figure 8**).

In the metacentric EAS2 (**Figure 4-E**) and EAS17 (**Supplementary Figure 8**) chromosomes, the centromeric function is embedded within a 37cen array, flanked by very short arrays of other repeats, including CENPB-sat regions of less than 10 kb. At EAS2, the CENP-A enriched regions correspond to the portion of the array with the highest sequence identity (**Figure 4-E**). At EAS17, the 37cen array is approximately 500 kb long and is entirely covered by the CENP-A binding domains in all the four individuals. Donkey C shows additional CENP-A enrichment peaks on single-copy sequences, suggesting the presence of rearrangements of this locus in the population (**Supplementary Figure 8**). No CENP-B binding was detected at these regions. In the metacentric EAS1 chromosome, a 50 Mb satellite array, primarily composed of 37cen repeats, was found (**Supplementary Figure 8**). This extensive array contains two main CENP-A enrichment peaks spaced 20 Mb apart, highlighting the complexity of this highly extended satellite region. No CENP-B binding was detected in this region.

In the metacentric EAS3 chromosome (**Figure 4-F**), the CENP-A enrichment peaks of the four individuals cover a 120 kb array of CENPB-sat. This is the longest CENPB-sat array in the donkey genome and unique in that it is bound by CENP-B (**Figure 4-F**, **Supplementary Figure 8**, **Supplementary Table 4**). Interestingly, EAS3, which represents the ancestral configuration present in the hypothetical common ancestor of equids, is orthologous to ECA2q and ECA3q[31], and ECA2 is the only horse chromosome where CENP-A and CENP-B colocalize (**Figure 4-C**). These findings reinforce our previously proposed model [17], suggesting that CENPB-sat represented the ancestral centromeric satellite whose centromeric function was subsequently lost at all loci except EAS3 and ECA2, which retain the ancestral organization.

In the remaining chromosomes (EAS6, EAS15, EAS20, EAS21, EAS22, EAS23, EAS24, EAS25, EAS26, EAS28 and EAS29), CENP-A binds the SatC satellite (**Figure 4-G**, **Supplementary Figure 8**). Similar to what was described in the other satellite-based centromeres, the satellite units are highly conserved in the CENP-A binding region. While SatC is the sole satellite in the centromeric region of the metacentric chromosomes EAS6 and EAS15, the acrocentric chromosomes (EAS20, EAS21, EAS22, EAS23, EAS24, EAS25, EAS26, EAS28 and EAS29) show a SatC centromeric region embedded within megabase-sized pericentromeric arrays primarily composed of SatA satellite (**Figure 4-G**, **Supplementary Figure 8**). The amount CENPB-sat at these centromeres is negligible with only very short stretches of a few kilobases and CENP-B binding is absent (**Figure 4-G**, **Supplementary Figure 8**). The positional variation at donkey satellite-based centromeres is less pronounced than in horses (**Figure 4-E-G**, **Supplementary Figure 8**). A possible explanation is that the high identity and homogeneity of centromeric satellite arrays mask the positional variation of CENP-A binding domains.

In agreement with our previous FISH results[7], in the donkey genome we detected megabase-sized satellite loci at non-centromeric terminal positions (EAS1pter, EAS4pter, EAS6pter, EAS9pter, EAS10pter, EAS11pter, EAS13pter, EAS18qter, EAS19qter, EAS27qter and EAS30qter) and in the q arm of EASX (**Supplementary Figure 9**). These non-centromeric loci contain different satellite families with variable levels of sequence identity. Short stretches of satellite repeats are otherwise present in other genomic positions, likely representing evolutionary traces of genomic rearrangements (**Supplementary Table 4**).

At EAS4pter, EAS6pter, EAS9pter, EAS10pter, EAS13pter, EAS18qter, EAS19qter, EAS27qter and EAS30qter, we detected arrays of the newly described 2PI subfamily - 2PI-telo (**Supplementary Figure 9, Supplementary Table 4**). Previous FISH experiments with telomeric probes showed intense signals on a few unidentified donkey chromosome ends[32,33]. FISH, using a probe for TTAGGG repeats, strong signals at EAS1pter, EAS4pter, EAS6pter, EAS9pter, EAS10pter, EAS11pter, EAS12pter, EAS13pter, EAS18qter, EAS19qter, EAS27qter, and EAS30qter (**Supplementary Figure 10**). With the exception of EAS12pter, this distribution overlaps with that of the 2PI-telo subfamily in the donkey genome (**Supplementary Figure 9, Supplementary Table 4**), indicating that these signals correspond to the 2PI-telo satellite. It is important to note that the donkey individual analyzed by FISH is not related to the mule used to assemble the donkey genome. Therefore, the additional EAS12pter signal detected by FISH may be attributable to population-level variation in the distribution of this satellite subfamily. On several chromosomes, the signal intensity differed between the two homologs (**Supplementary Figure 10**), further suggesting that polymorphism in the copy number of 2PI-telo repeats may be present in the population.

### Chromosome and centromere evolution in horse and donkey

The T2T horse and donkey genome assemblies represent a precious resource to investigate chromosome and centromere evolution in horse and donkey lineages. To this end, it is important to remember that the horse is considered the closest extant species to the equid ancestor and equid ancestral chromosomes generally correspond to horse whole chromosomes or their arms[31,34,35].

Thanks to the T2T assemblies a highly resolved chromosome alignment between horse and the donkey orthologs was carried out (**Supplementary Figure 11**). We confirmed chromosome orthologies that were previously described at the cytogenetic level but detected undescribed inversions at EAS4, EAS14, EAS23 and EAS25 and complex rearrangements at EAS17 (**Supplementary Figure 11**). These whole chromosome alignments allowed us to investigate chromosome evolution with respect to the position of centromeres and satellite DNA.

As shown in **Figure 5**, the majority of donkey chromosomes correspond to entire horse chromosomes or single horse chromosome arms, indicating that they did not arise from fusion. Chromosome fusions were detected only at EAS1, EAS4, EAS5, EAS7, EAS8 and EAS10. Differently, it is well described that EAS3, although corresponding to ECA2q/ECA3q, was not derived from a chromosomal fusion but represents the putative ancestral configuration[31,34].

**Figure 5:**
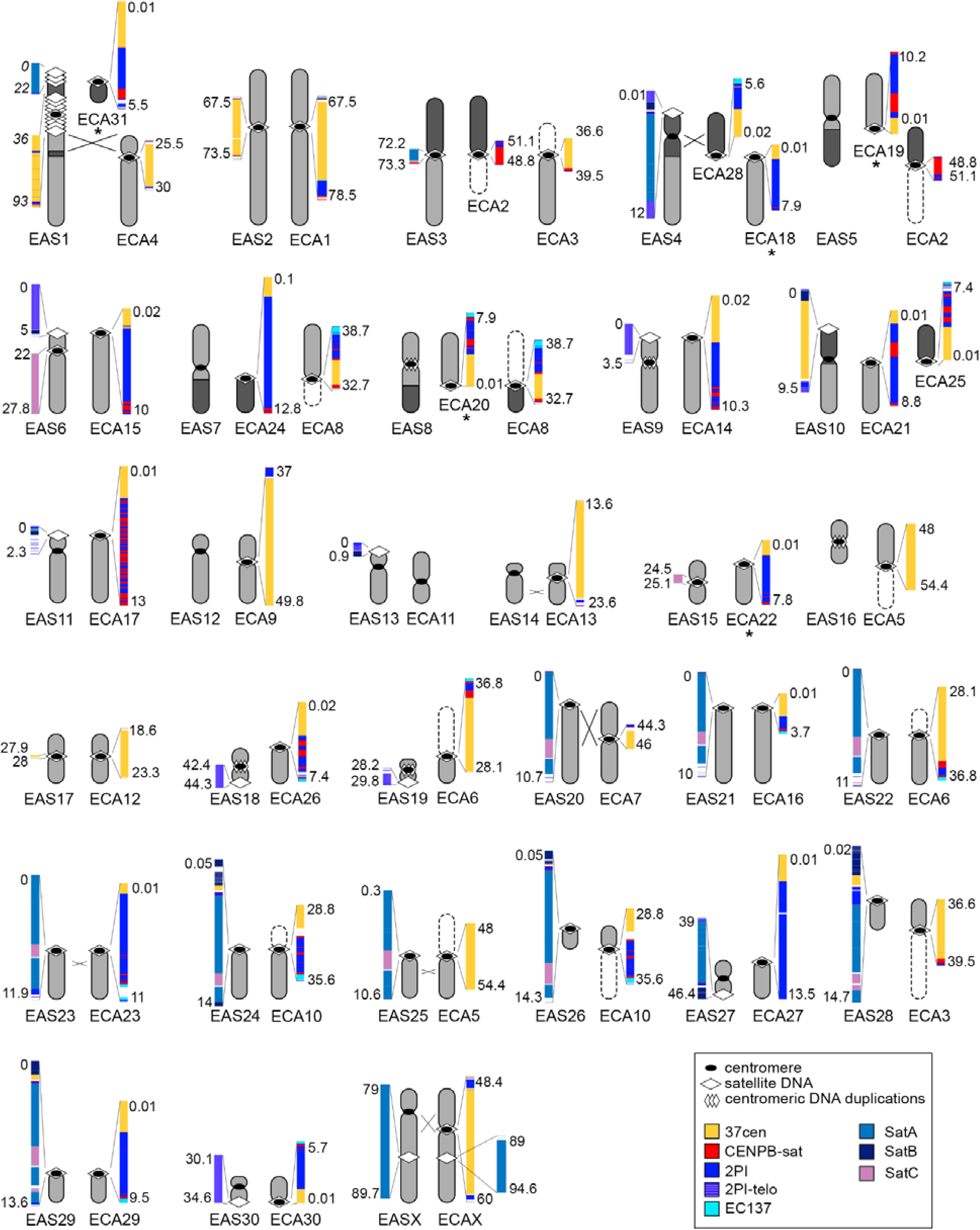
Satellite DNA and centromere organization in horse and donkey chromosomes. Donkey and horse orthologous chromosomes are shown, with centromeres (black ovals) and satellite DNA (white lozenges) indicated. For each satellite locus, a color-coded map of satellite DNA arrays is reported. Th lengths of satellite arrays are to scale, except for the large arrays at the p-terminus and peri/centromeric region of donkey chromosome 1. For horse chromosomes 18, 19, 20, 22, and 31 (marked with asterisks), the centromeric domains map to contigs anchored to the chromosome rather than to the primary scaffold.

With the exception of EAS1, all donkey chromosomes resulting from fusion carry satellite-free centromeres (**Figure 5**). However, the alignment dotplots between these donkey chromosomes and their horse orthologs show that these satellite-free centromeres are located several megabases away from the fusion points (**Supplementary Figure 11**). This suggests that the fusion may not be directly responsible for the formation of these satellite-free centromeres but rather that they originated through centromere repositioning events.The satellite-free centromere of EAS4 is located at an inversion breakpoint, suggesting that the newly detected EAS4 inversion may have played a direct role in its formation (**Supplementary Figure 11**). A similar pattern was observed in EASX, where the satellite-free centromere is positioned very close to an inversion breakpoint (**Supplementary Figure 11**). All the remaining donkey satellite-free centromeres (EAS11, EAS12, EAS13, EAS14, EAS16, EAS18, EAS19, EAS27 and EAS30) clearly originated from centromere repositioning events, consistent with our previous findings[8,19] (Carbone et al. 2006; Nergadze et al. 2018).

Comparison of horse and donkey orthologs revealed significant variations in satellite DNA composition, highlighting lineage-specific evolution of satellite families (**Figure 5**). However, in most cases, the chromosomal position of satellite loci is conserved among the two species. In particular, looking at **Figure 5**, we can identify six different situations: (1) maintenance of satellite DNA at horse and donkey orthologous peri/centromeric positions (EAS1cen/ECA4cen, EAS2cen/ECA1cen, EAS3cen/ECA3cen-ECA2cen, EAS17cen/ECA12cen, EAS21cen/ECA16cen, EAS22cen/ECA6cen, EAS23cen/ECA23cen, EAS24cen/ECA10cen, EAS25cen/ECA5cen and EAS29cen/ECA29cen). At EAS1cen/ECA4cen, EAS2cen/ECA1cen, EAS3cen/ECA3cen-ECA2cen andEAS17cen/ECA12cen, the same satellite DNA families are present in both species suggesting that these satellite loci remained conserved during evolution; (2) maintenance of satellite DNA at horse and donkey orthologous non-centromeric positions (EASXq/ECAXq) which share extended SatA arrays; (3) maintenance of satellite DNA at donkey non-centromeric terminal positions and horse orthologous centromeric positions (EAS1pter/ECA31cen, EAS6pter/ECA15cen, EAS9pter/ECA14cen, EAS11pter/ECA17cen, and EAS30cen/ECA30cen). The terminal donkey positions can be interpreted as remnants of ancient centromeres that were inactivated in the donkey and conserved in the horse; (4) presence of satellite DNA in the donkey centromeric or terminal positions and absence at horse orthologous positions (EAS6cen/ECA15, EAS15cen/ECA22, EAS13pter/ECA11). These loci correspond to repositioned donkey centromeres that acquired satellite DNA during evolution (EAS6cen and EAS15cen) or remnants of ancestral inactivated centromeres (EAS13pter); (5) loss of satellite DNA in the donkey compared to horse orthologous positions (EAS4/ECA28cen-ECA18, EAS5/ECA19cen-ECA2cen, EAS7/ECA24cen-ECA8cen, EAS8/ECA20cen-ECA8cen, EAS10/ECA21cen-ECA25cen, EAS12/ECA9cen, EAS14/ECA13cen, EAS15/ECA22cen, EAS16/ECA5cen and EASX/ECAXcen) resulting from chromosomal rearrangements or repositioning. By comparing the alignment dotplot coordinates in **Supplementary Figure 11** with the genomic coordinates of the short satellite repeat stretches identified by RepeatMasker in the donkey genome, we found that short arrays of different satellite families are present at the fusion regions of EAS1, EAS5, EAS7, EAS8, and EAS10, at an inversion breakpoint of EASX and in correspondence to horse satellite arrays at EAS12, EAS14, EAS15, EAS17 and EAS18 (**Supplementary Table 4**). These may represent remnants of the satellite arrays from the ancestral chromosomes; (6) presence of satellite DNA at terminal donkey positions and at the opposite centromeric end in the horse orthologous chromosome (EAS4pter/ECA28cen, EAS10pter/ECA25cen, EAS18qter/ECA26cen, EAS19qter/ECA6cen and EAS27qter/ECA10cen). This peculiar comparative localization can be interpreted as a consequence of satellite DNA exchange between opposite chromosomal termini[17,36,37].

### Annotation of transposable elements

RepeatMasker using Dfam 3.8 revealed an additional 124.6 Mb of transposable element derived sequence in EquAss-T2T_v2 compared to ASM1607732v2, and 23.9 Mb in TB-T2T compared to EquCab3.0. In EquAss-T2T_v2, all four major TE classes showed increases in content relative to ASM1607732v2: LINE elements (+78.7 Mb), LTR retrotransposons (+29.3 Mb), SINE elements (+15.1 Mb), and DNA transposons (+1.5 Mb). TB-T2T showed corresponding increases in content relative to EquCab3.0: LINE (+18.4 Mb), SINE (+2.6 Mb), LTR (+2.4 Mb), and DNA transposons (+0.5 Mb) (**Figure 6A**). Element counts similarly increased proportionally across all of the four major TE classes in both assemblies **(Figure 6B)**. A reduction of interspersed TE content of 42.3% to 39.0% in TB-T2T (vs EquCab3.0) and 42.4% to 41.4% in EquAssT2T_v2 (vs ASM1607732v2) was observed (**Supplementary Table 5**).

**Figure 6:**
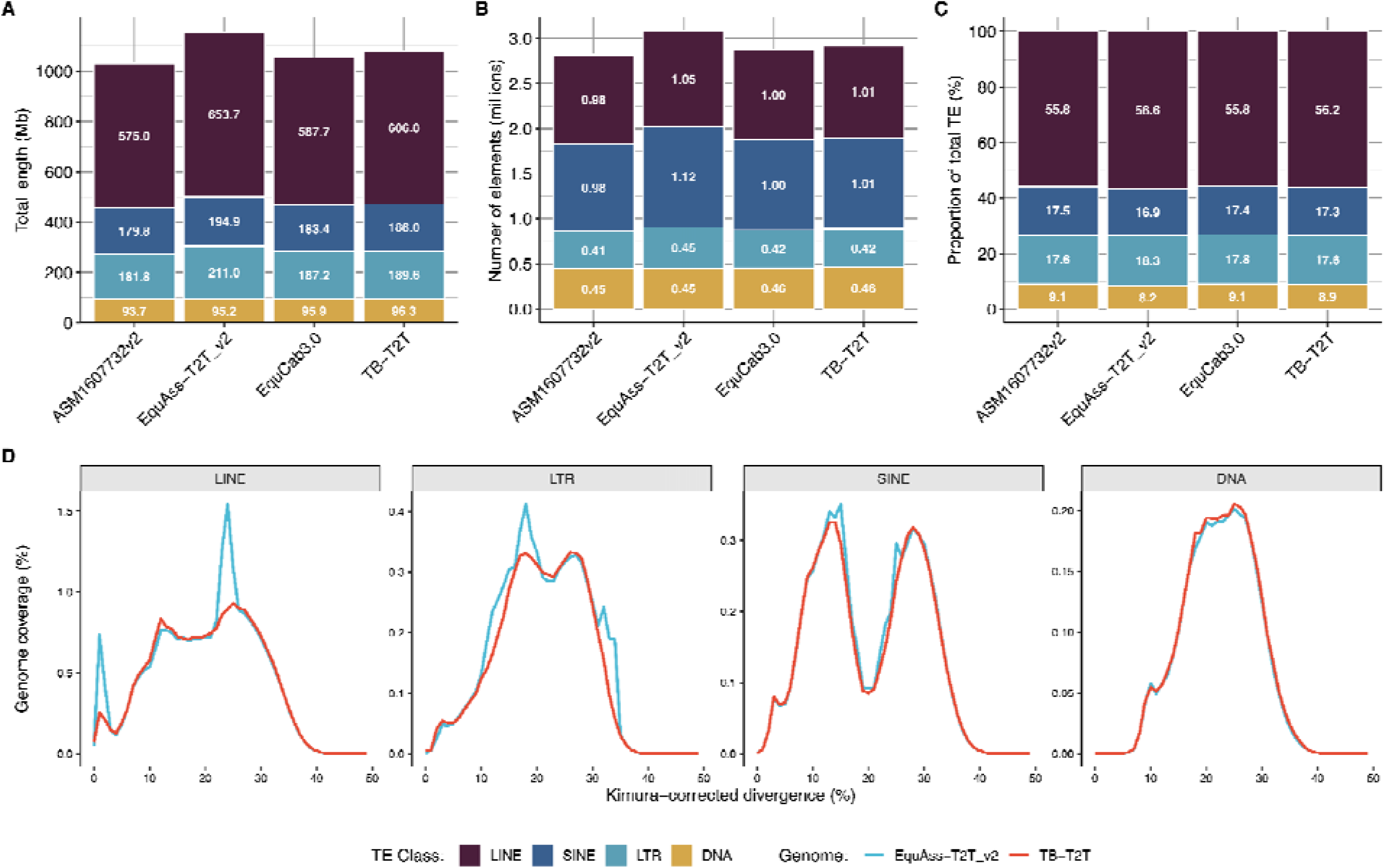
Transposable element composition of equid assemblies. **(A)** total base pairs per TE class for each assembly; **(B)** number of elements for each assembly; and **(C)** per-class composition proportional to total TE content per genome; for the four major transposable element classes (LINE: Long INterspersed Elements; SINE: Short INterspersed Elements; LTR: Long Terminal Repeats; DNA transposons). **(D)** Kimura-corrected divergence plot of LINE, LTR, SINE, and DNA transposable elements, showing the TE divergence distribution of both EquAss-T2T_v2 and TB-T2T. The SINE peak from 22-33% divergence represents activity of the older SINE/MIR elements, with SINE/tRNA activity dominating the more recent peak between 7-18% divergence. For LTR and LINE classes, EquAss-T2T_v2 expansion occurs at multiple regions of the divergence spectrum: for LTR, at 11-15%, 17-19%, and 32-35%; for LINE, at 1-3% and 23-25%. Plot resolution using 1% divergence bins.

Kimura divergence analysis showed the additional LTR and LINE element coverage in EquAss-T2T_v2 compared to TB-T2T is concentrated in relatively narrow divergence regions (**Figure 6D**).

### IsoSeq mapping and Tissue repository

IsoSeq data was generated for 13 tissues from this mule to benefit annotation efforts, and provides another opportunity to assess completeness. Ideally the IsoSeq measurement would catalog all transcripts for each of the tissues sequenced. If the genome is complete, all transcripts generated by the IsoSeq data would map to the assembly. This analysis is complicated by the fact that these transcripts would be produced from a pair of haplotypes that are genetically diverged from one another by 4 million years of evolution[28,29]. To account for this, the IsoSeq reads were mapped to the “Mule” assembly that contained both the TB-T2T, and EquAss_T2T_v2 chromosomes and contigs. Total reads were counted from the full length, non-concatemer files and are reported in **Supplementary Table 6**. The count of reads uniquely mapped to each haplotype were counted, and the number of unmapped reads were tallied for each tissue type as well. The raw datasets have been loaded to the SRA.

As a result of the success of the Equine FAANG Adopt a Tissue[38] effort, 133 tissues were harvested from this mule and are banked at Texas A&M for subsequent analysis. In most cases, 8 aliquots of each tissue type were collected and snap frozen in liquid N_2_. A full catalog of these tissues is provided as **Supplementary Table 7**. As described previously, 14 tissues were analyzed using PacBio Kinnex IsoSeq to benefit our initial annotation work. The remaining tissues are available to interested collaborators for a broad scope of genomic/epigenetic studies with the only condition being that of public release of the data.

### Equine high density SNP Chip

We mapped all probes from the EquCab3.0-based (GCF_002863925.1) Axiom MNEc670 SNP array to the TB-T2T genome by generating probe sequences from the array annotation file (Axiom_MNEc670 Annotation r3; ThermoFisher). The flanking sequence of each probe was expanded to include both alleles and formatted as a FASTA file, and aligned using blastn (BLAST+ 2.15.0)[39]. For each probe, the single best genomic hit was retained based on percent identity, alignment length, bit score, and e-value, and used the SNP position within the probe to compute its lifted coordinate on the T2T assembly.

Across the 670,796 probes in the array, 669,626 (99.8%) produced at least one BLAST alignment to the T2T genome. Of the probes with BLAST alignments, 608,444 (90.8%) mapped uniquely to a single genomic location, while 61,182 (9.1%) had multiple hits. Only 1,170 probes (0.17%) failed to map entirely under these alignment settings. Overall sequence identity was high, with 96.0% (642,585/669,626) of probes matching perfectly (100% identity) and 99.4% (665,335/669,626) mapping with at least 98% identity. Most BLAST aligned probes (627,677/669,626; 93.7%) retained their chromosome assignment between EquCab3 and the T2T genome. The lifted SNP positions were compiled into a probe-level annotation table and BED file, enabling translation of legacy SNP array genotypes from EquCab3.

## Discussion

Telomere-to-telomere assemblies are the essential next step for researchers working to understand the genetic bases of health and wellbeing, phenotypic diversity, and genomic divergence resulting from speciation. The TB-T2T, and EquAss-T2T_v2 assemblies presented here are the first step forward for the genomic scientists studying equids. Current efforts are underway to construct a pangenome for horses to enumerate structural variants across breeds, to annotate immune response genes, and to catalog short variants found across horse breeds. This will not be the last of these assemblies for the horse or other equids. It is our hope that the methods and resources presented here provide a foundation that will benefit efforts to understand centromeric, and repeat structure in those new assemblies.

The generation of telomere-to-telomere assemblies for the horse and the donkey has made it possible to draw a complete picture of the organization of both satellite-free and satellite-based centromeres in these two species. In the horse, satellite-based centromeres are typically made by an extended array of the 37cen satellite, which is the major centromeric satellite, flanked by other satellite families. Similar to what has been described for human centromeres[41–44], the functional horse centromeric domain is confined to an inner region of the 37cen array, which exhibits the highest sequence conservation compared to the pericentromeric portion of the array. In the donkey, where only half of the centromeres are satellite-based, the situation is much more heterogeneous, with multiple centromeric satellite families and significant variability in pericentric satellites. In donkey chromosomes, the functional centromere can indeed reside on a 37cen array (EAS1, EAS2 and EAS17), a CENPB-sat array (EAS3) or on a SatC array (EAS6, EAS15, EAS20-EAS26, EAS28 and EAS29).

The satellite-free centromeres of equid species allowed us to discover that the position of the CENP-A binding domains is not fixed in the population but slides in an about 600 kb window, giving rise to epialleles for CENP-A binding[8,12,16,18]. Here, the analysis of additional donkey individuals showed that CENPA binding domains can move within regions of up to 2.8 Mb, indicating that these genomic regions can be very large. Previous cytogenetic findings in horse spermatocytes suggested the presence of epiallelism for the position of the CENP-A binding domains at satellite-based centromeres[12]. Thanks to the telomere-to-telomere assemblies, we were able to confirm that centromere sliding occurs also at satellite-based centromeres in this species. This finding is consistent with the positional variation in centromeric domains reported at human and mouse satellite-based centromeres, indicating that this phenomenon is common in mammals[45–47].

The mapping of CENP-A and CENP-B ChIP-seq data on the horse and donkey T2T references confirmed the recent discovery that, in equid species, CENP-A and CENP-B domains are dissociated[17]. In our recent work, using a combination of cytogenetic and molecular approaches, we demonstrated that, in the horse, the satellite containing the CENP-B binding site (CENPB-sat) is found at a subset of pericentromeres and, at several loci, became degenerated losing the ability to be bound by CENP-B[17]. Using the horse T2T assembly, we were able to analyze each individual centromere at the sequence level that was not possible with the previous reference genomes. CENPB-sat arrays are present in only a subset of pericentromeres, and among these, only a fraction are bound by CENP-B. CENPB-sat arrays are typically intermingled with 2PI arrays at the periphery of the megabase-sized 37cen arrays. This organization supports our previous hypothesis that CENPB-sat and 2PI are ancestral satellites that were displaced in the pericentromeric area following the expansion of the 37cen satellite in the centromeric core[17]. This is consistent with the well-established models for satellite DNA expansion and evolution[48–51]. Our previous findings showed that the uncoupling between CENP-B and CENP-A is extreme in the donkey, where, although a functional protein is expressed, no CENP-B binding could be detected by immunofluorescence and CENPB-sat is very poorly represented in the genome, with only a unique CENPB-sat signal detected by FISH at the primary constriction of EAS3[17]. Analyzing the sequence of the T2T donkey assembly, we discovered that the longest CENPB-sat array(120 kb) was present on the EAS3 centromere. Only a portion of this array is bound by CENP-B, supporting evidence of satellite degeneration and explaining why we were not able to detect CENP-B signals by immunofluorescence[17]. At this centromere, the entire CENPB-sat array corresponds to the functional centromere, bound by CENP-A, similar to what was observed at the centromere of horse chromosome ECA2. Therefore,ECA2 and EAS3 centromeres are the only horse and donkey centromeres that retained the ancestral configuration, in which the CENPB-sat satellite was the centromeric satellite binding both CENP-A and CENP-B. These observations support our hypothesis that the uncoupling of CENP-A and CENP-B during equid evolution may have played a role in the extensive evolutionary reshuffling of equid karyotypes[17].

The comparison of horse and donkey telomere-to-telomere genomes made it possible to shed light on chromosome, centromere and satellite DNA evolution in horse and donkey lineages at the sequence level. The horse karyotype is considered the closest to the ancestral equid karyotype with a prevalence of acrocentric chromosomes. In the horse, the acrocentric chromosomes typically display long arrays of CENPB-sat and 2PI satellites flanking a short 37cen array, while most metacentric chromosomes carry long 37cen arrays with very short 2PI and/or CENPB-sat stretches at the periphery. This arrangement may be due to the relatively recent emergence of horse metacentric chromosomes through fusion, inversion, and centromere repositioning events[17,22,31]. These processes may have contributed to the reduction or loss of CENPB-sat and 2PI satellites, as well as the expansion of 37cen.

The donkey karyotype was restructured by numerous centromere repositioning events and chromosomal rearrangements[8,19,20,31,35]. These events occurred in very short and recent evolutionary times[28,31,34], explaining the presence of such a high number of satellite-free centromeres as well as the numerous non-centromeric satellite loci that represent remnants of ancestral inactivated centromeres. We confirmed that the majority of donkey satellite-free centromeres emerged following centromere repositioning. However, thanks to this new genome assembly, we discovered that two of them are located at an inversion breakpoint. A similar situation was previously described for a satellite-free centromere of Burchell’s zebra[15], indicating that inversions represent another mechanism for the emergence of satellite-free centromeres in equid species.

The analysis of the donkey chromosomes showed the expansion of satellite families which are absent or poorly represented in the horse genome such as SatA and SatC. An intriguing finding was that, at most donkey satellite-based centromeres, long SatA arrays are present in the pericentromeric region. This satellite family is composed of 4,678 bp monomers[17] which contain portions of identity with LINE L1-4B elements. It is well described that transposable elements can originate satellite repeats[52–55] and we previously demonstrated that LINE L1 elements are enriched at most equid satellite-free centromeres[8,9], and as such it is tempting to speculate that they may have triggered the expansion of satellite repeats. In addition, SatA monomers contain a region with high sequence similarity to the *Parascaris equorum* ETSTY7-like repeat, which was previously shown to be extensively amplified on the equid X chromosome and on several autosomes[56]. Taken together, these observations suggest that this sequence - proposed to have been horizontally transferred from *Parascaris* - expanded through the proliferation of SatA families. In addition, we discovered a donkey-specific expansion of the newly discovered 2PI subfamily (2P1-telo) which is composed of monomers made by telomeric repeats and 2PI repeats. This satellite subfamily is found at terminal non-centromeric positions, which could represent remnants of ancestral inactivated centromeres or the result of satellite DNA exchange between opposite chromosomal termini. This discovery confirmed a previous hypothesis, based on cytogenetic observations, that in this species a degenerate telomere-like satellite sequence could represent an abundant satellite family associated with heterochromatin[32,33]. The presence of satellite sequences containing telomeric repeats was described in other mammalian species, such as whales[57] and carnivores[58–60]. Similar to the donkey situation, these peculiar satellites were mainly localized at chromosomal termini[58–60]. Given the content in TTAGGG repeats of 2PI-telo repeats, sufficient to allow the identification of these satellite loci by FISH using a telomeric probe, and their chromosomal localization, we can properly classify them as heterochromatic interstitial telomeric sequences (het-ITSs). Het-ITSs were defined as extended blocks of telomeric-like repeats located at pericentromeric, terminal or intrachromosomal regions, generally coinciding with C-bands and easily detectable by cytogenetic analysis[61]. These sequences were identified in several vertebrate species, some insects and plants[61–68] and seem to be intrinsically prone to breakage[69–71]. Interestingly, most donkey 2PI-telo loci likely derived from satellite DNA exchange between opposite termini, supporting the fact that these sequences are prone to rearrangement.

Despite the lineage-specific variation in satellite DNA families, we identified several satellite loci that are present at conserved positions in both species. While the composition of satellite families is conserved at some loci, in several cases different satellite families expanded in the two lineages.In a few cases, these satellite loci underwent further expansion during donkey evolution, making them more extended than their horse counterparts. The most notable example is the very extended satellite arrays at the p terminus and at the pericentromeric region of EAS1, which are 22 Mb and 57 Mb long, respectively. These regions were previously identified at the cytogenetic level by FISH and were found to be polymorphic in size in the donkey population[72], highlighting the variability of satellite DNA organization among individuals.

The T2T assemblies revealed substantial TE increases (124.6 Mb in EquAss-T2T_v2, 23.9 Mb in TB-T2T) despite a proportional decrease of interspersed TE content from 42.3% to 39.0% in TB-T2T (vs EquCab3.0) and 42.4% to 41.4% in EquAssT2T_v2 (vs ASM1607732v2). This dilution effect reflects that the newly resolved regions are primarily composed of telomeric and satellite-rich centromeric sequences rather than TE-rich regions. LTR and LINE element expansions account for the majority of the additional TE content observed in EquAss-T2T_v2 relative to TB-T2T (**Figure 6A**). Kimura divergence analysis localizes these expansions to specific divergence ranges (**Figure 6D**), including ERV1, DIRS, and ERVK families within the divergence range spanning 11-35% (data not shown). Such patterns would traditionally be attributed to lineage-specific transposition activity; however, with T2T assemblies resolving large repeat-rich regions previously absent from prior assemblies, these patterns could also reflect segmental duplications or other chromosomal rearrangements. These patterns suggest the expansions may be largely attributable to segmental duplication, especially at higher divergence ranges, while the LINE elements at 1-3% divergence represent more likely candidates for recent insertional activity. Resolving the relative contributions of these mechanisms requires further study. While we hypothesize segmental duplication as the predominant mechanism, lineage-specific TE variation over comparable timescales has been documented in closely related taxa (e.g., Calidris sandpipers[73]), indicating some level of transposition-driven expansion remains plausible. More precise LTR age estimation requires analysis of rigorously verified LTR pairs, which the RepeatMasker annotation methods used here do not reliably identify. Such analysis would better distinguish lineage-specific transposition from segmental duplication and represents an avenue for future study.

Finally, the continued improvement of both long read sequencing technologies, and genome assembly algorithms coupled with ever improving computational power, and decreasing sequencing costs have served to democratize the work of genome assembly. Manual curation, and quality assurance remain domains for experts, but with this expertise genomes can be articulated very accurately and almost completely but for repeat regions whose lengths exceed that of contemporary ultra-long read technologies. The improvements made in the two phased haploid assemblies for the horse and donkey presented here are substantial in terms of increased composition of the autosomal and sex chromosomes (11.4% for the horse, and 11.5% for the donkey) over their corresponding reference assemblies they are replacing. The result is that accurate, sequence based analyses of centromeres and repeat regions is now possible that can inform our understanding of the chromosomal evolution events that created the equid species we have today.

## Methods

### Sample Collection

A healthy 20 year old female mule from Cornell University was euthanized under IACUC 1986-0216. Tissues were harvested immediately and flash frozen in liquid nitrogen. They were subsequently moved to Texas A&M University for long term storage. A list of the tissues can be found in Supplementary Table 7, and they can be requested by contacting the authors.

### Karyotyping

#### Isolation of mesenchymal stem cells

Equine peripheral blood MSCs were isolated from heparinized whole blood samples as previously described[74]. In short, peripheral blood mononuclear cells (PBMCs) were isolated using density gradient centrifugation and seeded in a culture flask with medium to promote MSC growth. After several days in culture, immune cells died off, and MSCs developed in adherent colonies, which were expanded and cryopreserved for use in experiments.

#### Cell cultures and chromosome preparations

Frozen mesenchymal stem cells of the female mule, were kindly provided by collaborators at Cornell University, Ithaca, NY, USA. Cell cultures and harvest for chromosome preparations were done following standard protocols[75,76]. Briefly, the cells were grown in alpha MEM with nucleosides and Glutamax (Gibco), supplemented with 20% FBS (Gemini Bioproducts) at 5% CO_2_. Semi-confluent (∼60-70%) and actively proliferating cultures were harvested with demecolcine solution (10 µg/mL; Sigma Aldrich), treated with Optimal Hypotonic Solution (Rainbow Scientific), and fixed in 3:1 methanol/acetic acid. The cells were dropped on clean, wet glass slides and checked under phase contrast microscope (x200) for quality.

#### Karyotyping and cytogenetic analysis

Chromosomes were stained by GTG-banding[77] for karyotyping. Images of 20 metaphase spreads were captured and analyzed using Axioplan2 microscope (Carl Zeiss, Inc., Jena, Germany) and IKAROS (MetaSystems GmbH, Altlussheim, Germany) software. The horse and donkey chromosomes were identified and arranged into karyotypes following the International System of Cytogenetic Nomenclature of the domestic Horse[78] and GTG-banded chromosome nomenclature of the donkey[32], respectively.

### Genome assembly

#### Genome Survey

To ensure high-quality Illumina sequencing data, we performed trimming and quality assessment as follows: We used Trim Galore (version 0.6.6) to preprocess the raw mule Illumina whole genome shotgun sequencing data. Trim Galore, which is based on Cutadapt (version 3.5. Cutadapt removes adapter sequences from high-throughput sequencing reads), removes low-quality bases, sequencing adapters, and other redundant sequences. The trimming was performed with the following parameters: A quality threshold of 20 was applied to remove bases with lower quality scores. Reads shorter than 20 bp after trimming and their mates were discarded to ensure the effectiveness of downstream analyses.

Following trimming, we assessed the data quality using FastQC (version v0.12.1, https://www.bioinformatics.babraham.ac.uk/projects/fastqc/). FastQC provides detailed quality reports, including base quality distribution, GC content, and sequence length distribution. This quality control step was crucial for verifying the accuracy of the input data and minimizing potential biases in downstream analyses.

We utilized Meryl[79] (version 1.3) to create a k-mer database from the cleaned Illumina sequencing data of the mule genome. k-mers of length 31 were selected, and Meryl was used to calculate the frequency of each k-mer in the dataset, resulting in a k-mer frequency histogram. The k-mer frequency histogram was then used as input for GenomeScope[80] 2.0, which applies a k-mer-based statistical approach to estimate genome heterozygosity(1.23%), repeat content(0.697), and genome size(haploid= 2,757,299,182 bp) from sequencing reads (**Figure 7**).

**Figure 7:**
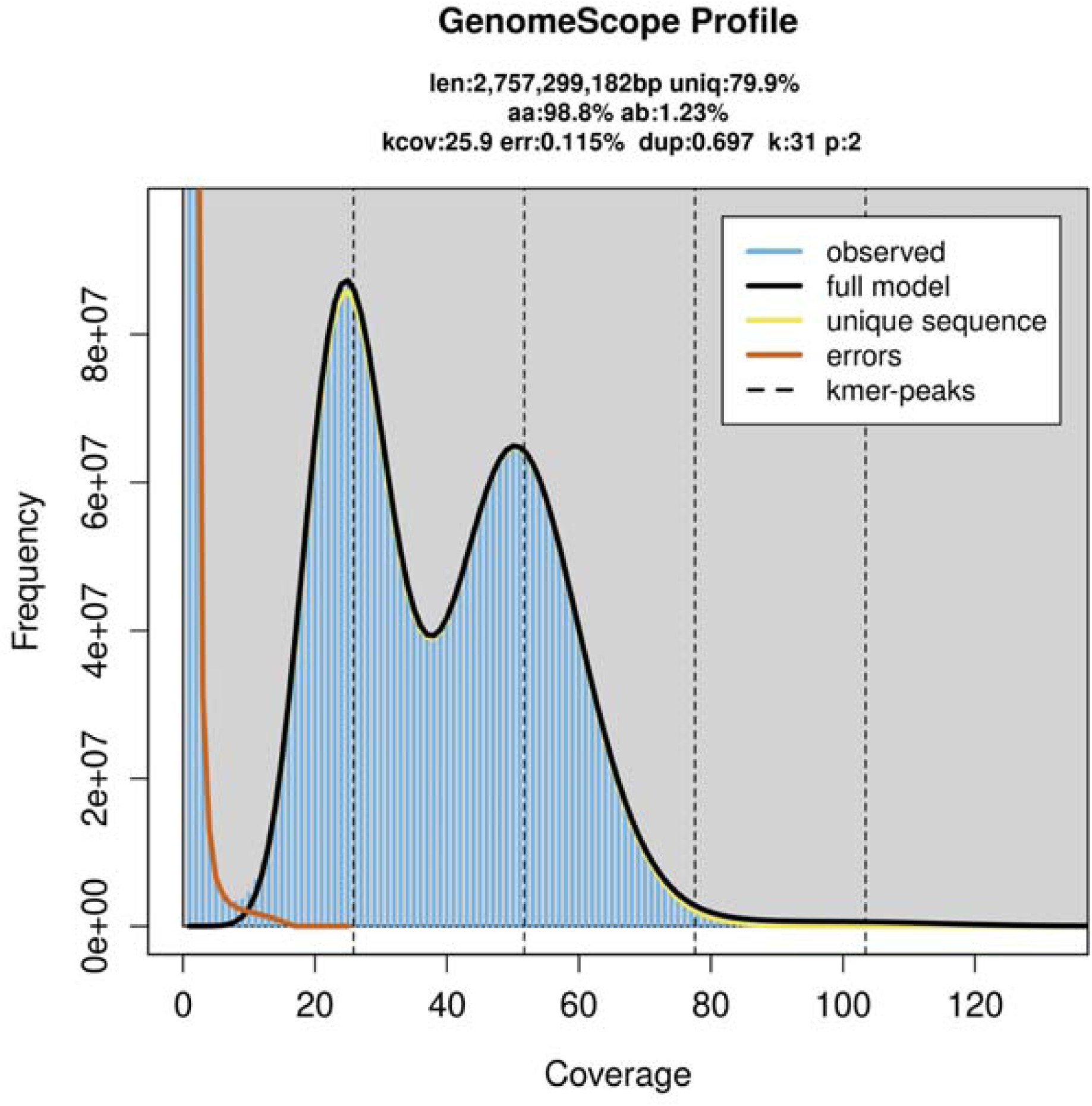
GenomeScope k-mer profile of the mule genome. GenomeScope analysis based on 31-mer frequency distribution estimated the mule genome size at approximately 2.76 Gb with 79.9% unique sequence content and a heterozygosity rate of 1.23%. The two major peaks represent heterozygous and homozygous k-mer coverage, respectively, while the fitted model (black line) closely follows the observed distribution (blue bars). The low estimated sequencing error rate (0.115%) and moderate duplication level (0.697) indicate high data quality suitable for accurate genome assembly.

#### Contig level assembly

For the initial assembly phase, we used the Hi-C mode of Verkko[81] (version 1.4.1) to construct the mule genome. Verkko’s Hi-C mode integrates multiple types of sequencing data to enhance assembly accuracy and contiguity. The input data included high-fidelity (HiFi) long reads from PacBio, Oxford Nanopore Technology ultra long (UL-ONT) reads, and Dovetail Omni-C data, all derived from the mule genome. In this version of Verkko, Omni-C data is used exclusively for phasing within contigs. Consequently, despite the Omni-C sequencing data originating from Mule, we were unable to initially obtain distinguishable haplotype contigs for either the Thoroughbred Horse or the Donkey. The sequencing data coverage is reported in Supplementary Table 8.

#### Assembly polishing

We utilized Inspector software[82] to correct small-scale assembly errors (< 50 bp) by replacement and local re-assembly of regions identified with structural errors (≥ 50 bp). This resulted in a high-accuracy assembly of the mixed two haplotypes. Using second-generation sequencing data from both parents and the Genome Analysis Toolkit (GATK version 4.4.0.0)[83], we distinguished the horse and donkey genomes by aligning the horse dam, and donkey sire to the mule assembly and sorting horse vs. donkey based on the fewest homozygous differences. Contigs containing mitochondrial contamination were discarded based on the depth and length of the original data returned to assembly. Then two haplotype genomes were polished using Pilon with clean trimmed parental Illumina whole genome shotgun sequencing data[84]. The trimming process and tools are the same with raw mule Illumina data. The polished scaffolds underwent gap filling using ONT data with RagTag[85] and RFfiller[86]. The nearly chromosome-level scaffolds were finally curated using Hi-C data with PretextView software[87].

#### Mitochondrial genome assembly

Taking advantage of the long length and high accuracy of HiFi data, assembling the mitochondrial genome was straightforward, as many reads in the HiFi dataset easily span the entire mitochondrial genome. Assembly with MitoHiFi[88] begins by providing the NCBI Taxonomy ID (in this case, 9796), Mitochondria Reference Finder locates the most closely related reference mitochondrial genome in the NCBI database. It extracts HiFi reads aligned to this reference genome, removes reads exceeding a certain length threshold to avoid the influence of Nuclear Mitochondrial DNA Sequences, and then assembles the reads using Hifiasm[89]. The resulting contigs are circularized, redundant sequences are removed from the assembly, and the chrMT is annotated.

Mitochondrial reads were a component of the Illumina whole genome sequencing data of the Dam Thoroughbred Horse. Therefore, it is unwise to use the Pilon and the horse’s whole genome sequencing data to polish the horse genome before the assembly and isolation of the mitochondrial reads. This may introduce over-polishing of the NUMT region. Thus, we recommend that researchers using the same assembly process should perform mitochondrial assembly and eliminate reads aligned to the mitochondrial genome before the polishing steps.

### IsoSeq mapping and evaluation

14 full length non-chimeric, clustered IsoSeq datasets from banked mule tissues (spleen, pituitary, lung, cerebellar vermis, PBMC, long bone marrow, loin adipose, heart left ventricle, heart left atrium, lamina front, adrenal cortex, kidney cortex, hypothalamus, and esophagus) were collected to aid in annotation and transcription of this assembly. The “Mule” genome was created from concatenating the TB-T2T, and EquAss_T2T_v2 chromosomes and contigs together to encompass both haploid assemblies. IsoSeq mapping was conducted using minimap2 version 2.22-r1101[90] with the following parameters: “-I 8g -ax splice:hq -uf” as indicated by the minimap guide. “-I 8g” was utilized to indicate an index batch size of equal to 8 GB which controls memory usage during indexing and was required due to the large size of the “Mule” genome. The parameters “-ax splice:hq -uf” indicate a spliced alignment preset for high quality IsoSeq reads and the input is full length nonchimeric reads. After the reads were mapped and aligned using minimap2, creating a bam file, SAMTOOLS version 1.13[91] was used to sort the alignments for downstream analysis and index the newly created bam file.

For evaluation of our IsoSeq mapped dataset, we decided to indicate the number of mapped and unmapped reads for each tissue. Read counts were conducted utilizing SAMTOOLS version 1.20 to account for bitwise flags. The parameter “samtools view file.bam -F 4 -F 256 -F 2048” noted the number of reads that mapped for the IsoSeq dataset whereas “samtools view file.bam -f 4” was used to count the number of reads that do not map and align to combined “Mule” assembly for our IsoSeq dataset.

### Centromere and satellite DNA annotation

#### Cell lines

Primary fibroblast cell lines from horse and donkey were previously described[7,17–19]. The cells were cultured in high-glucose DMEM medium supplemented with 20% fetal bovine serum, 2 mM L-glutamine, 1% penicillin/streptomycin and 2% non-essential amino acids. Cells were maintained at 37 °C in a humidified atmosphere of 5% CO2.

#### Identification and analysis of satellite DNA loci

Identification of satellite DNA loci was performed with RepeatMasker (Galaxy Version 4.0.9) available at the Galaxy web platform (Community 2022) using a library made by the consensus sequences of horse and donkey satellite DNA families[17,21,22]. The monomer size and the consensus sequence of each satellite are reported in **Supplementary Table 1**. Given the high sequence degeneration of the 2PI satellite, multiple consensus sequences of varying lengths were used. Satellite regions of each family located within 3000 bp were merged using Bedtools v2.30.0. Satellite sequences were annotated only in the chromosomal sequences. For the horse TB-T2T assembly, the unplaced contigs NW_027222394.1, NW_027222395.1, NW_027222392.1, NW_027222391.1, NW_027222343.1 were also included in the analysis. Telomeric repeats were identified by searching the (TTAGGG) 4 query by BLASTN using - evalue 10 -word_size 11 -gapopen 5 -gapextend 2 -reward 2 -penalty -3 -outfmt 7 parameters. Resulting matches located within 250 bp of each other were merged into single loci using Bedtools v2.30.0. For each satellite DNA region, sequence identity heatmaps were generated using ModDotPlot[92] with --bin-freq mode.

#### ChIP-seq with anti-CENP-A antibody and downstream bioinformatic analysis

For Horse G, Donkey C and Donkey D, chromatin from primary fibroblasts was cross-linked with 1% formaldehyde, extracted and sonicated to obtain DNA fragments ranging from 200 to 800 bp. Immunoprecipitation was performed as previously described[8] (Nergadze et al. 2018) using an anti-CENP-A serum[12]. Paired-end sequencing was performed with Illumina HiSeq2500 platform by IGA Technology Services (Udine, Italy). CENP-A ChIP-seq reads from the other individuals were previously described[8,16,17] and deposited in NCBI SRA Archive: Twilight (SRX19696555, SRX19696554), AH1 (SRX19696520, SRX19696519), AH2 (SRX19696543, SRX19696529), AH3 (SRX19696552, SRX19696551), AH4 (SRX19696550, SRX19696536), Horse A (SRX2789324, SRX2789325), Horse C (SRX2789347, SRX2789336), Horse D (SRX2789370, SRX2789369), Horse S (SRX23002276, SRX23002277), Donkey A (SRX2789326, SRX2789371), Donkey B (SRX2789329, SRX2789328). CENP-B ChIP-seq reads were previously described (Cappelletti et al. 2025) and deposited in NCBI SRA Archive: Horse C (SRX22972993, SRX22972992), Donkey B (SRX22972995, SRX22972994). ChIP-seq reads were aligned with Bowtie2 (version 2.4.2) using paired-end mode and default parameters[93]. Normalized enrichment peaks were obtained with the bamCompare tool available in the deepTools suite (3.5.0 version)[94] using RPKM normalization in subtractive mode. Plots were obtained with pyGenomeTracks (3.6 version)[95].

#### Fluorescence In Situ Hybridization

Metaphase spreads were obtained with the standard air-drying procedure. The telomeric probe, a mixture of 1-20 kb-long synthetic (TTAGGG)n fragments was prepared as previously described[96,97]. Hybridization to donkey metaphase spreads was carried out as previously described[98]. Post-hybridization washes were performed in low stringency conditions at 37 °C in 4x SSC 25 % formamide, 0.1 % Tween20 in 4× SSC. Chromosomes were counterstained with DAPI. Digital grey-scale images were acquired with a fluorescence microscope (Zeiss Axio Scope.A1) equipped with a cooled CCD camera (Photometrics) Pseudo-coloring and merging of images were performed using the IpLab software. Chromosomes were identified by computer-generated reverse DAPI banding according to the published karyotypes.

#### Comparative genomic analysis

Pairwise alignments between whole horse and donkey orthologous chromosomes were performed with Chromeister (version 1.5a)[99] using default parameters.

#### Repeat annotation

Repeat annotation was performed using RepeatMasker v4.1.7-p1 in species mode (-species Equus) with the sensitive flag enabled (-s) and configured to use the Dfam 3.8 full root partition as the library (dfam38_full.0.h5, 72GB uncompressed). Kimura-corrected divergence values for transposable element classes (LINE, SINE, LTR, DNA transposons) were obtained from RepeatMasker .align output files. Divergence values were calculated using the Kimura 2-parameter model with CpG correction to account for hypermutability at CpG dinucleotide sites, and binned at 1% divergence intervals from 0-50% using the parseRM Perl script[100].

#### Web access of source and derived datasets

All WGS, HiC, and IsoSeq datasets for the mule have been deposited in the SRA under the Bioproject PRJNA1136960. Illumina short read data for the donkey sire can be found at Bioproject PRJNA1135234, and for the Thoroughbred horse dam at Bioproject PRJNA1135235. Raw ChIP-seq data from this study are available in the NCBI SRA database under accession number PRJNA1354467.

## Acknowledgements

Funding support was provided by USDA/NIFA Blueprint Grant:2024-67015-42330. TK provided project leadership. KL and EC assembled and finished the genomes. EC and EG annotated centromeric elements of the genome, CD and DA annotated the repeat regions of the genomes. FP carried out molecular and cytogenetic measurements. JC, ER, LJ, NHAE, XA provided support, and QC analyses for the assemblies. MLS and EH generated the PacBio HiFi data. BDP, SK, BPW, SYB, CS, JCr, GB, AP provided the ONT UL data, assembly algorithms, and support for assembly curation. DA and DM, provided the mule for this project as well as parental DNA samples. JCu and BD provided SNP chip mapping data. JCu, SS, BD, MM, SDA, JLP, and TK are leading the Horse Pangenome Project. EC, EG, CD, KL, and TK wrote and edited the manuscript.

## Supplementary Figures

**Figure S1:** GTG-banded metaphase chromosomes of a female mule (63,XX). **(A)** Metaphase spread; **(B)** Autosomes of the horse; **(C)** Autosomes of the donkey, and **(D)** Horse and donkey X chromosomes

**Figure S2:** The horse ECA11 satellite-free centromere. The ChIP-seq profiles of CENP-A (orange, 10 individuals) and CENP-B (purple, 1 individual) are shown. The y-axis reports the normalized read counts whereas the x-axis reports the coordinates on the horse reference genome.

**Figure S3.** Horse satellite-based centromeres. For each centromere, the ChIP-seq profiles of CENP-A (orange, 10 individuals) and CENP-B (purple, 1 individual) are shown at the top. The y-axis reports the normalized read counts whereas the x-axis reports the coordinates on the horse reference genome. The colored bar represents the satellite DNA array, with colors denoting different satellite families as indicated in the legend shown in the last page. The sequence identity maps obtained with ModDotPlot are reported at the bottom. For chromosomes 18, 19, 20, 22, and 31 (marked with asterisks), the centromeric domains map to contigs anchored to the chromosome rather than to the primary scaffold (IDs: NC_091701.1, NC_091702.1, NC_091703.1, NC_091705.1, and NC_091714.1).

**Figure S4:** The non-centromeric satellite locus of horse chromosome X. The ChIP-seq profiles of CENP-A (orange, 10 individuals) and CENP-B (purple, 1 individual) are shown at the top. The y-axis reports the normalized read counts whereas the x-axis reports the coordinates on the horse reference genome. For Horse G, a low enrichment is present at SatA array and can be interpreted as a combination of mapping bias and peculiar satellite organization in this individual. The colored bar represents the satellite DNA array, with colors denoting different satellite families as indicated in the legend. The sequence identity maps obtained with ModDotPlot are reported at the bottom.

**Figure S5:** The 2PI-telo subfamily in donkey. Sequence of a 2PI-telo monomer. The sequence highlighted in yellow is made by telomeric repeats (red) and degenerated telomeric-like sequences. The blue part corresponds to 2PI repeats.

**Figure S6:** Donkey satellite-free centromeres without DNA duplications. The ChIP-seq profiles of CENP-A (orange, 4 individuals) and CENP-B (purple, 1 individual) are shown. The y-axis reports the normalized read counts whereas the x-axis reports the coordinates on the donkey reference genome.

**Figure S7:** Donkey satellite-free centromeres with DNA duplications. The ChIP-seq profiles of CENP-A (orange, 4 individuals) and CENP-B (purple, 1 individual) are shown at the top. The y-axis reports the normalized read counts whereas the x-axis reports the coordinates on the donkey reference genome. The sequence identity maps obtained with ModDotPlot are reported at the bottom.

**Figure S8:** Donkey satellite-based centromeres. For each centromere, the ChIP-seq profiles of CENP-A (orange, 4 individuals) and CENP-B (purple, 1 individual) are shown at the top. The y-axis reports the normalized read counts whereas the x-axis reports the coordinates on the donkey reference genome. The colored bar represents the satellite DNA array, with colors denoting different satellite families as indicated in the legend shown in the last page. The sequence identity maps obtained with ModDotPlot are reported at the bottom.

**Figure S9:** Donkey non-centromeric satellite DNA loci. For each region, the ChIP-seq profiles of CENP-A (orange, 4 individuals) and CENP-B (purple, 1 individual) are shown at the top. The y-axis reports the normalized read counts whereas the x-axis reports the coordinates on the donkey reference genome. The colored bar represents the satellite DNA array, with colors denoting different satellite families as indicated in the legend shown in the last page. The sequence identity maps obtained with ModDotPlot are reported at the bottom. Non-centromeric loci shorter than 500 kb are not represented here but reported in Supplementary Table 4.

**Figure S10:** FISH with a telomeric probe on donkey metaphase chromosomes. Donkey chromosome pairs with large terminal signals are indicated with their corresponding chromosome numbers. Because the exposure during image acquisition was calibrated on the strong signals of 2PI-telo loci, telomere signals appear very faint and are not visible on all chromosome ends.

**Figure S11:** Whole chromosome alignment plots between donkey chromosomes and horse orthologous chromosomes. The position of donkey satellite-free centromeres with and without DNA duplications are indicated with blue and red

## Supplementary Tables

**Table S1:** Summary of telomere lengths and residual assembly gaps in the TB_T2T and EquAss-T2T_v2 genomes. The table lists the lengths of telomeric regions at both chromosome arms (p and q) and the sizes of any remaining unassembled gaps in each chromosome. The results indicate that both assemblies are nearly complete, with telomeres identified at both chromosome ends and only a few minor residual gaps.

**Table S2:** Satellite families in horse and donkey. The different satellite DNA families of horse and donkey are listed. Monomer length and consensus sequence are indicated, together with the reference in which they were first described.

**Table S3:** Annotation of satellite loci in the horse TB-T2T genome. Coordinates of satellite arrays identified with RepeatMasker are provided for all chromosomal sequences and for the five contigs containing the p-terminal ends of chromosomes 18, 19, 20, 22, and 31 (NC_091701.1, NC_091702.1, NC_091703.1, NC_091705.1, and NC_091714.1).

**Table S4:** Annotation of satellite loci in the donkey EquAss-T2T_v2 genome. Coordinates of satellite arrays identified with RepeatMasker are provided for all chromosomal sequences.

**Table S5:** Transposable element annotation of TB-T2T, EquAss-T2T_v2, EquCab3.0, and ASM1607732v2 assemblies. Each row represents one TE subclass with element count, total base pair coverage, and genome percentage calculated using RepeatMasker utility buildSummary.pl. Element counts include all repeat fragments, including those contained within or overlapping other repeats, resulting in slightly higher counts (1-3% depending on TE class) than RepeatMasker’s standard .tbl output (used in Figure 6) which excludes overlapping fragments. For TB-T2T, EquAss-T2T_v2, and EquCab3.0, Kimura two-parameter and CpG-corrected divergence data are included as 50 bins of 1% increments (0-50%), indicating the base pair coverage within each divergence range. Summary table at bottom shows total counts and percentages per TE class per assembly, calculated from the subclass data above. Annotation performed using RepeatMasker with Dfam 3.8 and identical methods across all assemblies.

**Table S6:** Mapping of 14 full length non-chimeric, clustered IsoSeq datasets from banked mule tissues to the Mule assembly (created via combining two haplotypes of the horse and donkey). The reads were counted and filtered based on whether they mapped within the assembly using “samtools view”. The filter “samtools view -F 4 -F 256 -F 2048” was applied to count the reads that were mapped primarily based on bitwise flags. The filter “samtools view -f 4” was applied to count the reads that were unmapped by looking for reads based on bitwise flags.

**Table S7:** The list of tissues collected, and banked from the mule that produced the TB-T2T and EquAss-T2T_v2 assemblies. These tissues were flash frozen in liquid nitrogen, and are currently banked at Texas A&M University. The IsoSeq data described in the manuscript, and whose mappings are provided as supplementary data were derived from this resource. Please reach out to the corresponding author for access to these tissues.

**Table S8:** The sequence data, and their respective platforms and coverage used to assemble, polish, and QC the TB-T2T and EquAss-T2T_v2 genomes.

